# Systematic errors in enzymatic conversion limit cell-free DNA methylation specificity

**DOI:** 10.64898/2026.03.24.713040

**Authors:** Netanel Loyfer, Judith Magenheim, Alaa Darwish, Sara Isaac, Jay Ganbat, Hussam Babikir, Anjeet Jhutty, Jonathan Wan, Antoni Bayés-Genís, Elena Revuelta-López, Amir Eden, Ravi Solanki, Yuval Dor, Tommy Kaplan

## Abstract

Enzymatic methylation sequencing (EM-seq) converts unmethylated cytosines to uracils while preserving DNA integrity, making it attractive for liquid biopsies. Here we report a reproducible fragment-level over-conversion error in EM-seq, which is not observed in bisulfite-based conversion or in Oxford Nanopore sequencing. While bisulfite and nanopore errors occur at sporadic CpGs, EM-seq generates molecules that appear fully unmethylated, introducing a false-positive background signal that severely limits deconvolution specificity in cfDNA analysis.

## Main text

Whole-genome bisulfite sequencing (WGBS) is the gold standard for DNA methylation analysis^1–3^ but suffers from substantial DNA degradation, limiting its applicability in low-input samples and the potential for fragmentomics analysis. Enzymatic methylation sequencing (EM-seq) has emerged as a promising alternative that preserves DNA integrity and fragment length^4^, and is being widely adopted for applications such as cell-free DNA (cfDNA) methylome analysis^5^. EM-seq relies on protection of methylated cytosines by TET-mediated oxidation and glucosylation, followed by APOBEC-mediated deamination of unprotected cytosines^4^. In principle, this yields methylation information equivalent to WGBS while avoiding chemical damage.

Indeed, bulk methylation levels measured by EM-seq and WGBS are highly concordant across many genomic contexts. However, as cfDNA assays increasingly rely on single-molecule resolution^3,6–8^, the structure of technical noise becomes as important as its average magnitude.

We first compared EM-seq and WGBS in a fully methylated pUC19 plasmid. The patterns of cytosines read as thymidines (indicating unwanted deamination of methylated cytosines) were strikingly different: while WGBS had a random distribution of isolated thymidines, in EM-seq we often observed multiple thymidines in the same molecule (**Fig. 1A,B**).

**Figure 1.**
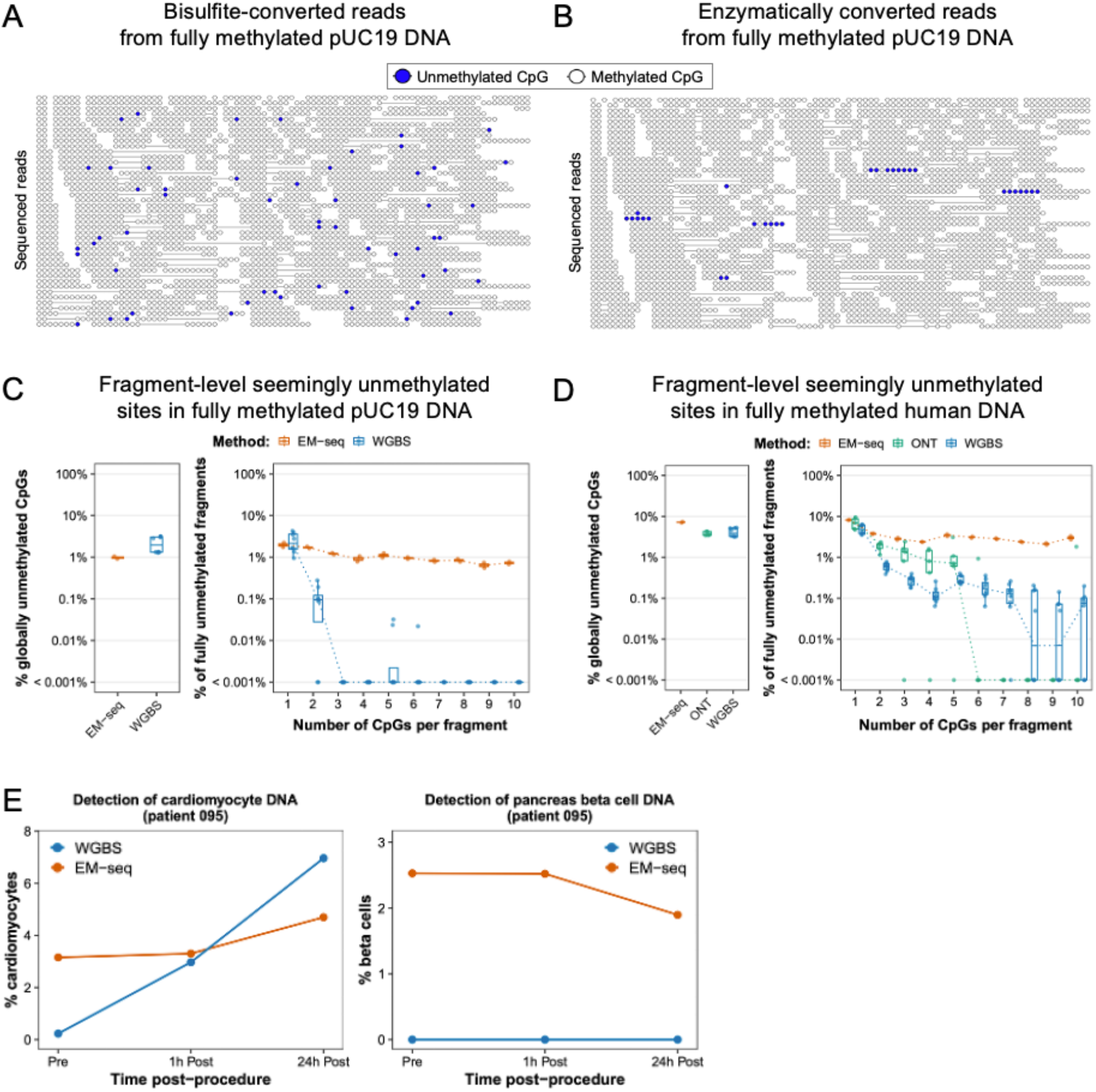
**A-B. Fragment-level over-conversion in EM-seq.** Representative pileup of multi-CpG sequenced reads at the pUC19 spike-in plasmid (position pUC19:1,155-2,652), taken from the same sample, sequenced by WGBS (A) and EM-seq (B). In bisulfite sequencing, unmethylated CpGs appear sporadically along reads, consistent with independent CpG-level noise. In EM-seq, a subset of reads appear fully unmethylated across multiple CpGs, indicating fragment-level over-conversion. Blue: unmethylated CpGs; white: methylated CpGs. While CpG-level demethylation in the bisulfite treated sample (2.8%) is higher when compared to EM-seq (0.9%), the rate of unmethylated fragments is much lower (0.003% vs. 0.56%). Shown are ∼300 randomly sampled fragments. **C. CpG-level and fragment-level error rates in EM-seq**. The abundance of fully unmethylated fragments remains high for EM-seq, even for fragments with multiple CpGs, demonstrating fragment-level errors in conversion. **D**. Similar analysis as in (C) for human DNA, focusing on 1068 ubiquitously hyper-methylated regions^3^, across 10 cfDNA samples. Data include EM-seq (Illumina), WGBS (Illumina and Ultima Genomics) and Oxford Nanopore (ONT), showing a decay in fully unmethylated reads in WGBS and ONT, but not in EM-seq. **E**. cfDNA deconvolution results illustrate spurious tissue attribution in EM-seq data driven by fragment-level over-conversion, compared with matched WGBS libraries. Shown are cardiomyocyte-derived cfDNA in samples from a patient before and after myocardial catheterization (left), and beta cell-derived cfDNA in the same samples, as negative control (right).

Overall, the conversion efficiency is high in both methods, with 1% apparently unmethylated sites in EM-seq, and 2% in WGBS. Yet, conditional on an unmethylated CpG, the probability that the next CpG is also unmethylated site is 2% in WGBS, comparable to the overall proportion of unmethylated sites; in EM-seq this probability is 72.5%, indicating strong within-fragment dependence. Accordingly, the abundance of fragments containing multiple unmethylated sites remained ∼1% in EM-seq but rapidly decays in WGBS, as expected from random independent noise (**Fig. 1C**).

To explore whether this phenomenon extended to human genomic DNA, we leveraged a reference set of 1,068 genomic regions we found to be fully methylated across all major human cell types^3^. In these constitutively methylated regions, any observation of unmethylated DNA must arise from technical conversion errors rather than biological heterogeneity or rare cell populations. We assessed methylation patterns across three platforms: a single normal cfDNA sample sequenced in 10 independent libraries (EM-seq, n=2; WGBS, n=8), and an additional 5 normal cfDNA samples sequenced using Oxford Nanopore Technology (ONT). As shown in **Fig. 1D**, the probability of observing a fully unmethylated fragment declined sharply with increasing CpG count, in both WGBS and ONT, but remained at 2.5% in EM-seq.

This distinction has critical implications. In WGBS and ONT, CpG-level noise is exponentially suppressed at the fragment level as CpG density increases, enabling highly specific inference from multi-CpG markers^3,7,8^. In EM-seq, fragment-level over-conversion events occur at similar frequency regardless of CpG count, imposing an irreducible background of fully unmethylated molecules.

This error structure is particularly problematic in cfDNA applications that detect rare unmethylated fragments at loci that are otherwise fully methylated in other cell types contributing to normal plasma^3,9–11^. In this setting, a single over-converted EM-seq fragment is indistinguishable from a genuinely unmethylated cfDNA fragment and generates false-positive signal in downstream deconvolution analyses.

To assess this impact, we performed atlas-based deconvolution using markers unmethylated in a given cell type^3^ and quantified the proportion of cardiomyocyte-derived cfDNA in plasma from a patient with myocardial infarction, undergoing cardiac catheterization. While WGBS produced the expected pattern of elevated cardiomyocyte cfDNA after the procedure, EM-seq detected a spurious unmethylated signal at baseline that masked procedure-related elevation (**Fig. 1E**). As a negative control, we assessed the presence of pancreatic beta cell cfDNA in these samples. Consistently, while WGBS detected no beta cell-specific hypomethylated markers, EM-seq detected a strong, clearly artefactual signal.

Given the use of datasets from multiple sources and 3 sequencing platforms, the observed effects cannot be explained by differences in sequencing depth, fragment length distributions, or library complexity. We observed the same fragment-level over-conversion pattern in additional publicly available EM-seq datasets. Although the biochemical origin of over-conversion in EM-seq remains to be resolved, the observed pattern is consistent with failed protection of methylated cytosines prior to APOBEC-mediated deamination, resulting in rare but complete hyper-conversion of individual DNA molecules.

We note that this fragment-level noise does not substantially affect bulk methylation averages, explaining why it may remain undetected in standard benchmarking analyses. However, for fragment-resolved applications such as cfDNA deconvolution, rare-cell detection, or single-molecule methylation analysis, it establishes a lower bound on achievable specificity. While computational correction strategies may partially mitigate this effect, they require explicit modelling of fragment-level errors and cannot fully restore the exponential noise suppression characteristic of WGBS.

In summary, although WGBS and EM-seq exhibit similar per-CpG error rates, they differ fundamentally in how technical noise propagates across DNA fragments. Our findings urge caution in the adoption of EM-seq for liquid biopsy studies and highlight the need for improved enzymatic protection or for methods that explicitly identify and account for fragment-level errors.

## Notes

### Competing Interest Statement

The authors have declared no competing interest.

## References

1. Lister, R. et al. Human DNA methylomes at base resolution show widespread epigenomic differences. Nature 462, 315–322 (2009).

2. Ziller, M. J. et al. Charting a dynamic DNA methylation landscape of the human genome. Nature 500, 477–481 (2013).

3. Loyfer, N. et al. A DNA methylation atlas of normal human cell types. Nature 613, 355–364 (2023).

4. Vaisvila, R. et al. Enzymatic methyl sequencing detects DNA methylation at single-base resolution from picograms of DNA. Genome Res. 31, 1280–1289 (2021).

5. Wang, Y. et al. Molecular evidence of multiorgan damage in patients with immune-related adverse events. N. Engl. J. Med. 393, 2377–2379 (2025).

6. Landau, D. A. et al. Locally disordered methylation forms the basis of intratumor methylome variation in chronic lymphocytic leukemia. Cancer Cell 26, 813–825 (2014).

7. Guo, S. et al. Identification of methylation haplotype blocks aids in deconvolution of heterogeneous tissue samples and tumor tissue-of-origin mapping from plasma DNA. Nat. Genet. 49, 635–642 (2017).

8. Onuchic, V. et al. Allele-specific epigenome maps reveal sequence-dependent stochastic switching at regulatory loci. Science 361, (2018).

9. Sun, K. et al. Plasma DNA tissue mapping by genome-wide methylation sequencing for noninvasive prenatal, cancer, and transplantation assessments. Proc. Natl. Acad. Sci. U. S. A. 112, E5503–12 (2015).

10. Lehmann-Werman, R. et al. Identification of tissue-specific cell death using methylation patterns of circulating DNA. Proc. Natl. Acad. Sci. U. S. A. 113, E1826–34 (2016).

11. Moss, J. et al. Comprehensive human cell-type methylation atlas reveals origins of circulating cell-free DNA in health and disease. Nat. Commun. 9, 5068 (2018).

