## Supplementary material for "Systematic errors in enzymatic conversion limit cell-free DNA methylation specificity": Methods

### Sample collection and DNA sources

Cell-free DNA (cfDNA) samples were obtained from healthy donors and from a patient undergoing myocardial infarction and cardiac catheterization. Plasma was processed using standard double-spin centrifugation protocols to minimize cellular contamination. cfDNA was extracted using silica-based methods optimized for the recovery of short fragments. Fully methylated control DNA (pUC19 plasmid) were spiked-in prior to library preparation.

### Library preparation and sequencing

For each cfDNA sample, multiple independent sequencing libraries were prepared using enzymatic methylation sequencing (EM-seq; New England Biolabs) and whole-genome bisulfite sequencing (WGBS), following manufacturer-recommended protocols. Figure 1A-D shows ten libraries generated from a single cfDNA sample, including EM-seq (n=2), WGBS with Illumina paired-end short-read sequencing (NovaSeq X Plus, 2×150 bp, n=4), and WGBS with Ultima Genomics single-end sequencing (UG100, ~350bp, n=4). We also used cfDNA samples processed using Oxford Nanopore Technologies (ONT, n=5) for direct methylation detection. An additional set of samples from an MI patient was collected independently (3 timepoints) and sequenced (Ultima Genomics UG100), comparing bisulfite and enzymatic conversions. Publicly available EM-seq datasets, as well as additional in-house generated WGBS datasets, were also incorporated for validation analyses (Supplemental Table S1).

### Read processing and alignment

For samples analyzed via the Ultima Genomics UG100 platform, read alignment and deduplication were performed using the proprietary Ultima Genomics Analysis Pipeline with its integrated, platform-specific aligner. Samples sequenced using Illumina platforms were aligned using bwa-meth[1]. Samples sequenced using the Oxford Nanopore Technology platform were processed using Dorado (v0.8.2, https://github.com/nanoporetech/dorado). All samples were aligned to the human reference genome hg38 merged with the spike-ins Lambda and pUC19. PCR duplicates were removed using coordinate-based deduplication (sambamba, v0.6.8)[2]. Only uniquely mapped reads with MAPQ≥10 were retained for further analyses.

### Methylation calling and fragment-level analysis

Raw reads (bam/cram) were analyzed using wgbstools[3,4] for fragment-level methylation calling. For Figure 1C-D, fragments were classified as “fully unmethylated” if ≥80% of the covered CpGs were called as unmethylated.
To assess dependence between consecutive CpGs within a fragment, we calculated the conditional probability that a CpG is called as unmethylated given that the preceding CpG (on the same fragment) was also called as unmethylated.

### Cross-platform comparison

To assess whether observed error patterns were platform-specific, we compared EM-seq, WGBS, and ONT data across matched genomic regions. For each platform, we computed the frequency of fully unmethylated fragments as a function of CpG density. Decay of error rates with increasing CpG count was used as an indicator of independence between CpG errors. Exponential decay is expected under independent noise (as observed in WGBS and ONT), whereas deviation from this behavior indicates correlated errors.

### Spike-in analysis (pUC19 control)

To assess technical conversion errors, we used the fully methylated pUC19 plasmid, and examined read-level methylation patterns. Figures 1A-B were visualized using wgbstools pat_fig[3,4]. We quantified both CpG-level error rates (fraction of CpGs incorrectly unmethylated), and fragment-level error rates (fraction of fragments appearing fully unmethylated). These analyses enabled direct comparison between stochastic CpG-level errors and fragment-level coordinated conversion events.

### Fully methylated genomic regions

To assess technical error in human genomic DNA, we used a reference set of 1,068 genomic regions identified as constitutively methylated across all major human cell types. Based on the genomic segmentation of Loyfer et al.[4], we selected regions with ≥4 CpGs and average methylation ≥93% that was seen in all cell types.

### cfDNA deconvolution analysis

For Figure 1E, the percent of cardiomyocyte-specific cell-free DNA fragments was estimated using an atlas-based deconvolution framework[4]. Briefly, genomic loci that are uniquely unmethylated in a specific cell type but methylated elsewhere were used as biomarkers (top 25 markers, with ≥4 CpGs per analyzed fragment). Cardiomyocyte-derived cfDNA was quantified in samples collected before and after cardiac catheterization. Pancreatic beta cell markers were used as a negative control. Comparisons between EM-seq and WGBS were performed using matched samples and identical marker sets to isolate the effect of conversion error structure.

### Supplemental Table S1

| Dataset | Type | Condition | Platform | Method | Subjects (n) | Samples (n) | pUC19 spike-in | Notes | Source |
|---|---|---|---|---|---|---|---|---|---|
| **Healthy cfDNA** | cfDNA | Healthy | Illumina NovaSeq X Plus + Ultima Genomics | EM-seq (v2) + WGBS | 1 | 10 | Yes | Replicates from a single sample: 2 EM-seq + 4 WGBS (Illumina) + 4 UMBS (Ultima) | Prima Mente |
| **Healthy cfDNA ONT** | cfDNA | Healthy | Oxford Nanopore PromethION | ONT (5mC + 5hmC) | 5 | 5 | No | 5 healthy; low coverage; no spike-in | Hebrew Univ. |
| **MI cfDNA** | cfDNA | Myocardial Infarction | Ultima Genomics UG100 | EM-seq (v1) + WGBS | 1 | 6 | Yes | Matched pairs; 3 serial timepoints (A, B, C) | CIBERCV Spain |
| **Alzheimer's cfDNA ONT** | cfDNA | Alzheimer's disease | Oxford Nanopore PromethION | ONT (5mC + 5hmC) | 5 | 5 | No | 5 AD; low coverage; no spike-in | Hebrew Univ. |
| **Healthy cfDNA** | cfDNA | Healthy | Ultima Genomics UG100 | EM-seq (v1) + WGBS | 5 | 13 | Yes | Matched pairs; 3 subjects have EM-seq replicates | Hebrew Univ. |
| **GRAIL blood** | cfDNA + gDNA | Healthy | Illumina NovaSeq 6000 | WGBS | 23 | 46 | No | Pairs of cfDNA + WBC | Loyfer et al.[4] |
| **Loyfer Atlas** | gDNA | Normal human cell types | Illumina NovaSeq 6000 | WGBS | 139 | 205 | No | 39 different cell types | Loyfer et al.[4] |
| **Alzheimer's cfDNA** | cfDNA | Alzheimer's disease | Illumina NovaSeq 6000 | EM-seq (v2) | 10 | 10 | Yes | 10 AD | Prima Mente |
| **Gowrisankar et al.** | cfDNA | Colorectal cancer + controls | Illumina NextSeq2000 | EM-seq (v1), hybrid capture | 220 | 236 | Yes | 68 colorectal cancer + 36 advanced adenoma + 29 benign polyps + 87 healthy controls | **Gowrisankar et al.**[5] |

### Supplemental Bibliography


1. Pedersen, B. S., Eyring, K., De, S., Yang, I. V. & Schwartz, D. A. Fast and accurate alignment of long bisulfite-seq reads. *arXiv [q-bio.GN]* (2014).
2. Tarasov, A., Vilella, A. J., Cuppen, E., Nijman, I. J. & Prins, P. Sambamba: fast processing of NGS alignment formats. *Bioinformatics* **31**, 2032–2034 (2015).
3. Loyfer, N., Rosenski, J. & Kaplan, T. wgbstools: a computational suite for DNA methylation sequencing data analysis. *Life Sci. Alliance* **9**, (2026).
4. Loyfer, N. *et al.* A DNA methylation atlas of normal human cell types. *Nature* **613**, 355–364 (2023).
5. Gowrisankar, S. *et al.* Extracellular Vesicle Gene Expression Enables Sensitive Detection of Colorectal Neoplasia. *Genomics* (2025).
